# Underlying beneficial effects of Rhubarb on constipation-induced inflammation, disorder of gut microbiome and metabolism

**DOI:** 10.1101/2022.06.24.497289

**Authors:** Han Gao, Chengwei He, Rongxuan Hua, Chen Liang, Boya Wang, Yixuan Du, Yuexin Guo, Lei Gao, Lucia Zhang, Hongwei Shang, Jingdong Xu

**Affiliations:** Department of Physiology and Pathophysiology, Basic Medical College, Capital Medical University, 100069, Beijing, China; Department of Clinical Medicine, Basic Medical College, Capital Medical University, 100069, Beijing,China; Eight Program of Clinical Medicine, Peking University Health Science Center, 2018, 100081, Beijing,China; Department of Oral Medicine, Basic Medical College, Capital Medical University, 100069, Beijing, China; Department of Biomedical Informatics, School of Biomedical Engineering, Capital Medical University, 100069, Beijing, China; Class of 2025, Loomis Chaffee School, 4 Batchelder Road, Windsor, CT 06095, USA; Experimental Center for Morphological Research Platform, Department of Physiology and Pathophysiology Basic Medical College, Capital Medical University, 100069, Beijing, China

**Keywords:** Constipation, Rhubarb extract, Gut microbiome, Polyamine, SCFA.

## Abstract

**Background:** Although constipation is a common syndrome and a worldwide health problem. Constipation patients are becoming younger with a 29.6% overall prevalence in the children, which has captured great attention because of its epigenetic rejuvenation and recurrent episodes. Despite the usage of rhubarb to relieve constipation, novel targets and genes involved in target-relevant pathways with remarkable functionalities should still be sought after.

**Materials and methods:** We established a reliable constipation model in C57B/6N male mice using intragastric administration diphenoxylate and the eligible subjects received 600mg/25g rhubarb extraction to ameliorate constipation. Resultant constipation was morphological and genetically compared with the specimen from different groups.

**Results:** The constipation mice exhibited thicker muscle layers, improved content of cytokines, including IL-17 and IL-23, and lower content of IL-22. The bacterial abundance and diversity varied tremendously. Notably, the alterations were reversed after rhubarb treatment. Additionally, SCFA and MLCFA were significantly influenced by constipation accompanied by enhanced expressions of SCFA receptors, GPR41 and GPR43.

**Conclusion:** This thesis has provided an insight that rhubarb promoted the flexibility of collagen fiber, reduced pro-inflammatory cytokines and enhanced anti-inflammatory cytokines, and maintained intestinal microflora balance with potential effects on affecting the metabolism of fatty acids and polyamines.

## 1. Introduction

Constipation is a common clinical symptom of gastrointestinal dysfunction with a high international average incidence rate of 15%. It is characterized by difficult or infrequent passage or hardness of stool, and/or a feeling of incomplete evacuation (1, 2). With the common lifestyle of high consumption of sugar and fat, the prevalence of constipation is estimated as 20% or higher. This has serious effects on the quality of people’s life regardless of age and gender (3). Rhubarb is an essential traditional Chinese medicinal herb that has been applied to clinical practice for relieving constipation. A majority of current research on constipation has focused on motility enhancement. Motility of the gastrointestinal tract is an imprecise term embracing several measurable phenomena, including enteric contractile activity, gut wall biomechanical functions, and intraluminal flow responsible for the propulsion of gut contents (4). In order to uncover whether rhubarb exerted influence on gut motility, we evaluated correlations between the changes on muscles and collagen fibers density to demonstrate the sensitivity enhancements of colonic contraction.

The vast majority of gut microbes represents an extremely complex microenvironment assembly of an estimated 10-100 trillion symbiotic bacteria per individual, which are present in intimate contact with the host and correlate with health and disease (5, 6). A number of studies provide strong evidence that the microbes and their hosts share a wide range of resources needed to support physiological requirements (7, 8). More importantly, the gut microbiota actively produces a deal of immune regulatory metabolites (9).

Short-chain fatty acids (SCFAs), major end products of gut microbial fermentation and an energy source of epithelial cells, take part in regulating the gut immune response (5, 10). They promote mucin production and the expression of antimicrobial peptides (11). G-protein-coupled (GPR) receptors, such as GPR41 and GPR43, serve as SCFA receptors and facilitate SCFA to activate multiple cells including epithelial cells, adipocytes, and phagocytes, as well as regulate diverse cellular functions (12). Additionally, there is evidence that SCFA and its receptors contribute to acute inflammatory responses in the intestine (13).

Biogenic amines are conventionally produced via microbial fermentation of undigested amino acids by deamination, deamination-decarboxylation or carboxylation (14, 15). To the best of our knowledge, most polyamines in this region of the colon are produced depend on intestinal flora; amino acids can serve as precursors for polyamine production (16, 17). Based on fecal sample analysis, naturally abundant polyamines include putrescine, spermidine, spermine and cadaverine in the human colon (18, 19). The putrescine, spermidine and cadaverine are derived from the decarboxylation of ornithine, methionine and lysine, respectively. Dysregulation of the level of polyamine and its amino acid precursors has been found to be connected with inflammation and autoimmune diseases (20). However, the underlying mechanism by which polyamine is possessed remains poorly uncovered.

Metagenomics has begun to study the composition and genetic potential of the gut microbiota to demonstrate the breadth of the functional and metabolic potential of microbes. There is no doubt that there exists a link between constipation and microbiota (21). However, there appears to be no existing data that proves what role the microbiota played in relieving constipation after treating rhubarb. Accordingly, we performed metagenomics to demonstrate significant metabolic discrepancies among the groups to find out the effect of constipation and rhubarb extract on microbiota. On a side note, the microbiota would be transacting more directly with the host immune system and metabolism in the intestinal epithelium. Conceivably, the microbiota probably would be more directly involved in inducing constipation.

However, current research has been descriptive in relieving constipation by increasing bowel motility. Apart from that, the underlying mechanism by which rhubarb extract is possessed remains poorly addressed. Therefore, this study makes a major contribution to research constipation by demonstrating its critical mechanism of understanding how the constitutive constipation response is regulated by rhubarb extract.

## 2. Results

### Results1 Effects on the Feeding Behavior and Stool Parameters

First, to establish whether diphenoxylate and rhubarb administration influences the feeding behavior and excretion parameters, the details about the number, weight, and water content of the fecal pellets are the most intuitive indexes to assess constipation under laboratory conditions (22). Therefore, alterations in food intake, water consumption, urine volume, and stool parameters were also measured daily in four groups mentioned above. As shown in Figure 1B, food intake was dramatically reduced after treatment with rhubarb or diphenoxylate, respectively in comparison to the control group and the impact was returned to near-normal levels by treating constipation mice with rhubarb, but no significant difference in water intake and urine volume among the various groups. While a slight bodyweight decrease was observed in constipation-induced by diphenoxylate, an enhancement after administering rhubarb was shown (Fig. 1C). To address this issue, the fecal pellets of excreted daily collected from metabolic cages were evidently decreased in the model group compared to the control group (Fig. lD, n=9, p<0.001), while the pellets enhanced with rhubarb were administered. Furthermore, varied fecal color and shape are essential measurements of constipation. As shown in Figure 1E, the feces were irregular in size and shape with variably gray color in the constipation group, while the pellets got washy or even unshaped after treatment with rhubarb. However, these classical studies confirmed a significant increase in the water content of feces in the rhubarb group (Fig.1D, n=9, p<0.01) compared to the pellets of feces (Fig.1D, n=9, p=0.7437). The feces in the constipation group had the least water content and were remarkably enhanced after being treated with rhubarb extract. Also, we observed the number of feces in the colon was remarkably reduced in the rhubarb group as compared to the control group as shown in Figure 1D (n=9, p<0.01), which may contribute to explaining the more general phenomenon in the constipation mice treatment with rhubarb. Next, we determined whether the functional defecation was accompanied by abnormal alterations of intestinal length. As Figure 1F indicated, the measurement of colon length from ileocecum to distal colon in each mouse showed significantly longer colons in all rhubarb-treated mice, regardless of in normal mice or in constipation mice(Fig.1D, n=9, p<0.01), but no clear differences was observed between constipation group and the control group. Overall, these results validated that the constipation models were successfully achieved and rhubarb extract had clearly evolved defecation benefit by partly enhancing the fecal water content. Changes in weight loss, fecal water content, and number of defecation granules are also sensitive and responsible for the phenotype caused by rhubarb extract.

**Figure 1.**
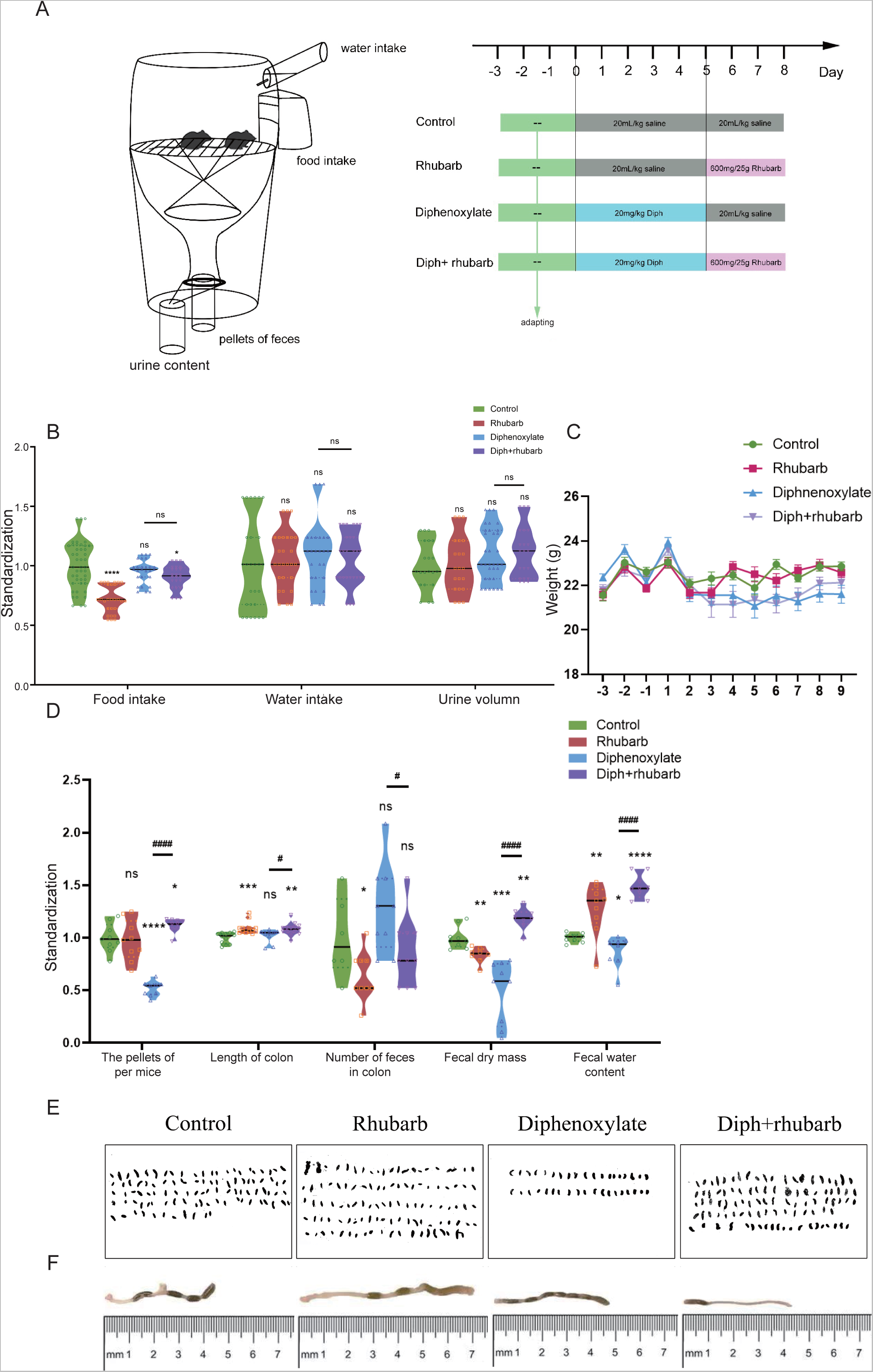
The experimental design and diphenoxylate-induced constipation model with its reinstatement of rhubarb treatment. (**A**) A timeline detailing the acclimatization, grouping, and time points for the drug treatments as well as the experimental paradigm. (**B**) Assessments of the consumption of food and water intake per day and urine volume per day. (**C**) Recording the body weight of mice fed with normal saline, rhubarb, diphenoxylate, and both diphenoxylate and rhubarb during the course of the experiment. (**D**) The violin graph represents the differences in the number of pellets defecated, fecal dry mass, fecal water content, and the length of colon section as well as the number of feces in the colon at the timeline of sacrifice in different groups. (**E**) Observation of the feces from one mouse per day to assess the feces characteristics in different groups. (**F**) Mouse colon photographs and their lengths as well as the number of feces in them are measured by the distance between anus (at 0) and cecum. All data from **C** and **D** is normalized by the control group. All values are presented as means ± SEM (n=9 per group). ns, P>0.05; *, P<0.05; **, P<0.01; ***, P<0.001. * vs control group; ^#^ vs diphenoxylate group.

### Results2 Alterations of histopathological and cytological structure of colon

We next investigated the associated changes in the histopathological and cytological structure of the colon induced by constipation and rhubarb intervention. We examined intestinal epithelial information by means of H&E staining (Fig. 2A). First of all, alterations in thicknesses of the colonic mucosa, submucosal, muscle layer were analyzed (23). The results showed that the layered muscle structure of the mouse colon under constipation status became thicker (Fig. 2B, n=9, p<0.001), which is markedly reduced after treatment constipation model with rhubarb extract (Fig. 2B, n=9, p<0.001). A contrary trend was detected in the thickness of the mucosa layer and rhubarb treatment induced the enhancement of mucosa layer (Fig. 2B, n=9, p<0.001). The thickness of the mucosa layer in constipation, whereas did not differ compared to the control group (Fig. 2B, n=9, p>0.05). There was a significant decrease in the thickness of the submucosal layer in the constipation group compared to the control group, and this trend was reversed by the treatment of rhubarb (Fig. 2B, n=9, p<0.001).

**Figure 2.**
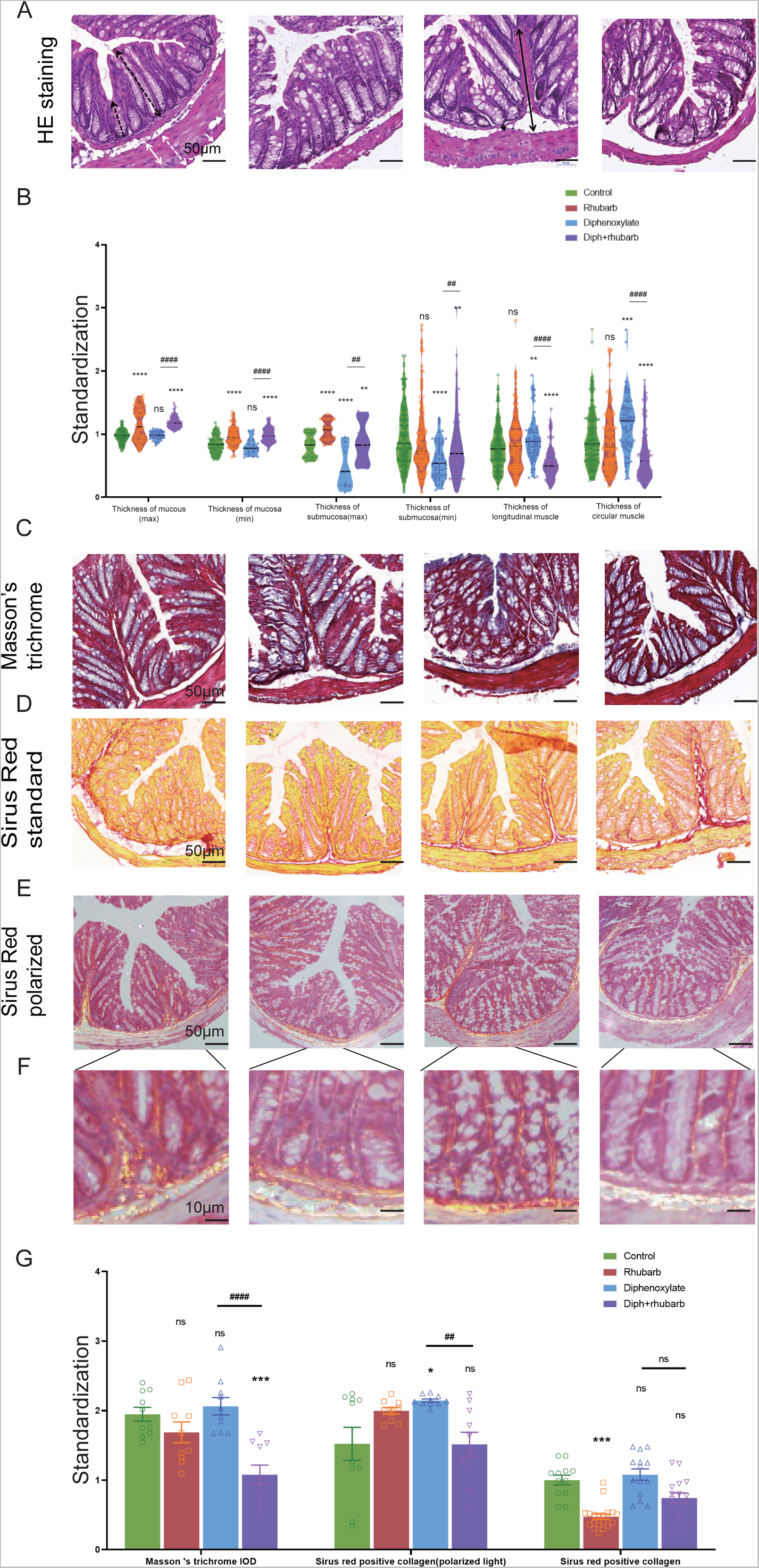
Histochemical colonic tissue stains show some routine qualitative differences for comparative purposes among four groups. (**A**) H & E staining reveals the general characteristics presented in a comprehensive overview. Black dotted line with double arrows: mucus layer (max). Black dotted line with single arrows: mucus layer (min). Black line with double arrows: submucosa layer (max). Black line with single arrows: submucosa layer (min). White dotted line with double arrows: muscle layer. White line with double arrows: circular muscle layer. White line with single arrows: longitudinal muscle layer. (**B**) The violin graph indicated the quantitative analysis of that the thickness of mucus layer (max and min), submucosa (max and min), and muscle layer (circular and longitudinal muscle). The data were normalized by the control group. (**C**) Representative images of Masson’s Trichrome staining to assess the fibrotic changes in the colon. (**D**) Total collagen in colon sections is stained with Sirus Red. Representative images are shown with the magnification of 10x. Representative picture of Sirus Red staining in polarized light, magnification, 10x. (**E**-**F**) Sirus Red-stained sections observed with polarized light, magnification 20x. Scale bars, 50μm. (**G**) The bar graph is then measured and quantified analysis of interstitial fibrosis as well as the total collagen amount by Sirus Red staining. The figures are successively from control group, rhubarb group, diphenoxylate group, and Diph+rhubarb group. All data is presented as means ± SEM (n=9 mice per group). ns, P>0.05; *, P<0.05; **, P<0.01; ***, P<0.001. * vs control group; ^#^ vs diphenoxylate group.

Considering that the muscle layer had a noticeable impact on the contraction, we conjectured that fibrosis might be an advanced-stage phenotype regulated by collagen fiber rather than an early causal factor in the development of hardness increases. In order to uncover whether fibrosis was involved in the process, all of these samples were observed by Masson’s trichrome and Sirius red staining. From Figure 2C, using Masson’s trichrome staining, we can see the constipation group contained more fiber, while it had less after rhubarb administration compared to the control group. In line with the results, what is interesting about the data in Figure 2D using Sirius Red staining was that quantification of collagen deposition showed a remarkably reduced following treatment with rhubarb. However, the polarized result revealed that the fiber was markedly increased, while the rhubarb group did not exhibit decreased tendency. To conclude, these data provided strong evidence that collagen fiber over-expression plays a causal role in increasing contraction intensity in muscle, which in turn, leads to a decrease in muscle strength.

To further validate these dominant effects on fiber, we assessed the strength and modulus of collagen fiber by means of AFM. The elastic moduli of smooth muscle, especially in the digestive tract, are still largely unexplored . An accurate mean modulus can be obtained only if the thickness of the colon tissue section is known . This value is based on the mean tissue rupture force and deformation of intestinal smooth muscle under fresh frozen sections. MLCT-BIO was chosen for the characterization. As Figure 3A-D showed the average elastic moduli of control, Rhubarb, Diphenoxylate-induced constipation, and constipation model treatment with Rhubarb measured by using MLCT-BIO. As shown in Figure.3B and E, the group in Rhubarb had the lowest modulus of 580.9 ± 111.4 KPa, while the group in constipation showed a much higher modulus of 4663 ± 305.2 KPa (Fig. 3C and E). Notably, as Figure 3D and E suggested, the moduli in the group of constipation treatment with Rhubarb showed a sharp decrease to 1396 ± 219.6 KPa (Fig. 3D and E). All these data were compared to the modulus in the control group. It was noted that all results of the modulus in constipation showed a significant increase, indicating that the elasticity of smooth muscle would also be significantly increased. This change, due to a physiological point of view, was consistent with the previous increase in collagen fibers in order to eliminate stool in the colon. Simultaneously it was also observed that after the use of rhubarb, the content of collagen fiber in the intestinal tissue was easily decreased due to the increase of moisture in the intestinal tract and the increase in movement speed, indicating that its elasticity would also be correspondingly increased. This change was consistent with the decrease in the content of collagen fiber measured in the previous experiment. Therefore, slight elastic changes in smooth muscle tissue caused by changes in collagen fibers can be assessed by atomic mechanics microscopy modulus.

**Figure 3.**
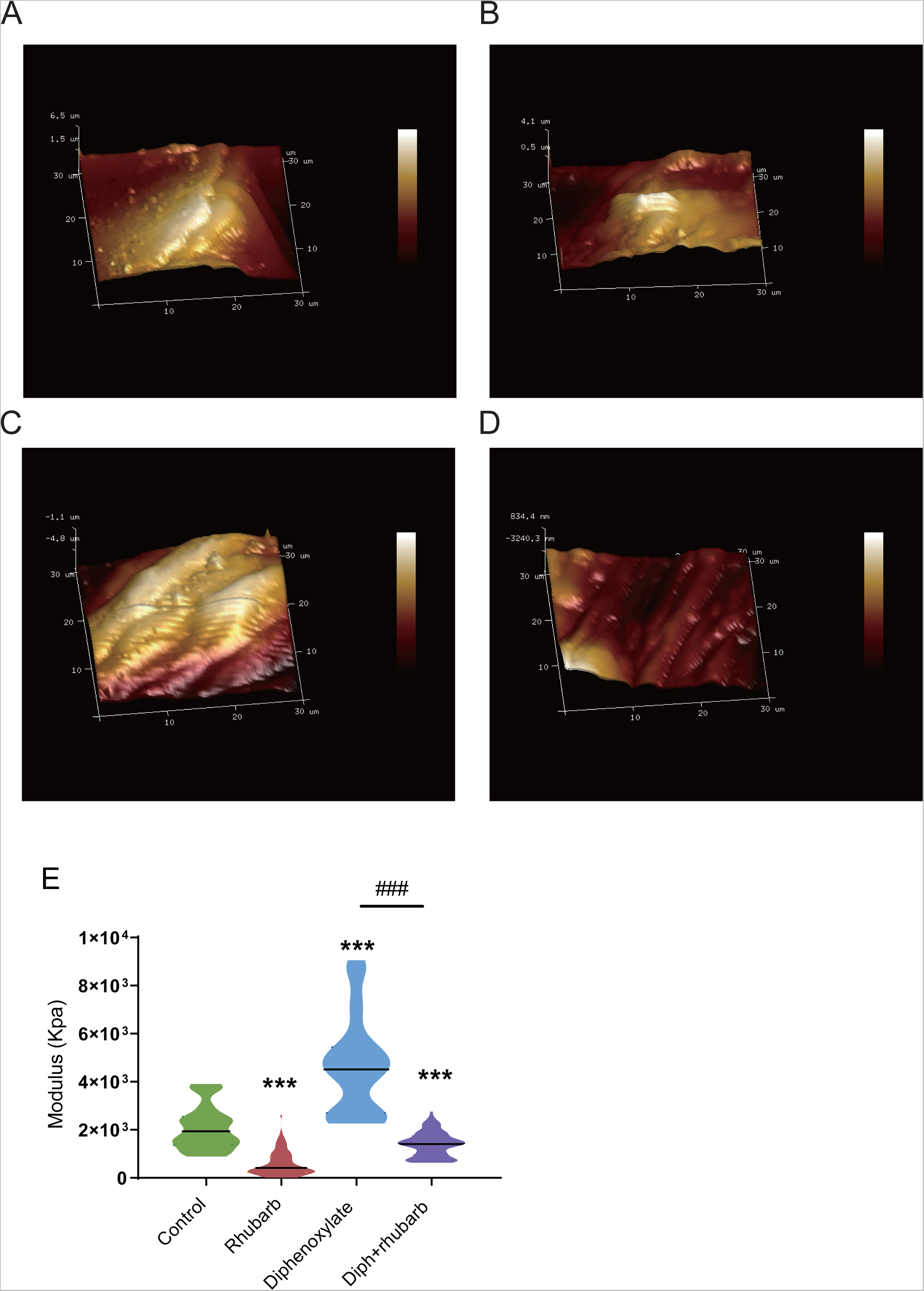
Atomic force microscopy (AFM) 3D images for the four groups: control group (**A**), rhubarb group (**B**), diphenoxylate group (**C**), and Diph+rhubarb group (**D**). (**E**) The violin graph indicates the modulus of the colon muscular from four groups. ns, P>0.05; *, P<0.05; **, P<0.01; ***, P<0.001. * vs control group; ^#^ vs diphenoxylate group.

### Results3 Measurement of cytokine concentrations

Colon crypt mucin has been found to be regulated by cytokines (24, 25). The change in the submucosa layer indicated that inflammation might be involved. To further elucidate the complex relationship between constipation and inflammation in intestinal epithelial cells, we next sought to determine the expressions of some cytokines such as IL-15, IL-17A, IL-22 and IL-23. IL-15, with pro-inflammatory effects, however, there was no difference among the four groups as Figure 4A indicated. IL-17 recruits neutrophils into the cecal mucosa to protect from the invasion of bacteria, but induce excessive inflammation (26). In alignment with our expectation, constipated mice predisposed to induce inflammation cause the level of IL-17A to be the highest. In addition, the high concentration decreased after treating constipation mice with rhubarb. Amongst the current research, the prevailing view is that IL-22 is mainly related to the maintenance of mucus barrier function by promoting LGR5^+^ epithelial stem cell regeneration/proliferation (27). And the results indicated that the IL-22 concentration in serum dropped in the constipation group and peaked in the Rhubarb group, which implied that rhubarb may play a protective role through increasing IL-22 level. IL-23 induces neutrophil polarization and promotes inflammation. As Figure 4A shows, we identified a noticeable decrease of IL-23 expression in the groups which were administered rhubarb regardless of the control mice or the constipation mice. In parallel to the above pro-inflammation cytokine, the constipation group had the highest level of IL-23.

**Figure 4.**
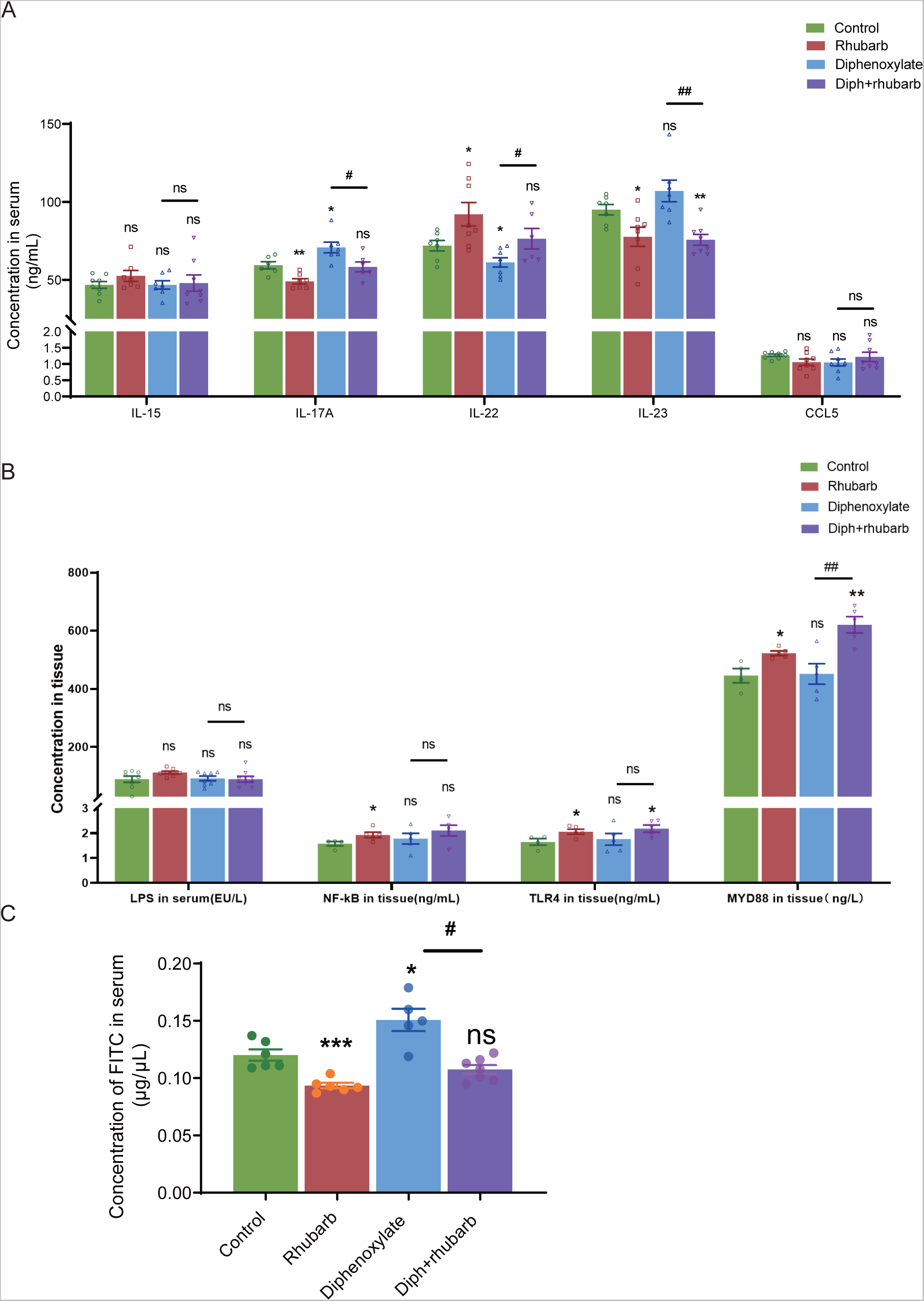
Measurement of the cytokine protein levels per cytokine concentrations and paracellular barrier function by paracellular tracer flux assays using FITC-dextran (150,000) among the four groups. (**A**) The bar graph illustrates the cytokine concentrations including IL-15, IL-17A, IL-22 and IL-23, CCL5 in the plasma. There is no obviously change as visualized distinct patterns in the IL-15 as well as CCL5 among the four groups. (**B**) The bar graph exhibits the concentrations of LPS, NF-κB, TLR4, and MyD88 in colonic tissue by ELISA among the four groups. (**C**) The concentrations of FITC-dextran in the circulating plasma in four groups are measured. The figure represents the combined results of repeated twice with similar results, and the results are expressed as the means ± SEM of six to eight mice per group. ns, P>0.05; *, P<0.05; **, P<0.01; ***, P<0.001. * vs control group; ^#^ vs diphenoxylate group.

To further determine the role of constipation on inflammation, we widened our search to screen for the concentration of lipopolysaccharide (LPS) in serum and its receptors, Toll-like receptor 4(TLR4) and myeloid differentiation primary response gene 88 (MyD88), in tissue (Fig. 4B). The results of nuclear factor-κB (NF-κB), TLR4, and MyD88 were coincident. More precisely, the level in the treatment with rhubarb group was particularly increased compared to the control group and dropped in the constipation group.

### Results 4 Impaired epithelial barrier function in constipation mice

The mucus layer is a vital physical barrier to both microbiota and toxin. For example, damage to gut barrier integrity, including the mucus layer, epithelial cell junctions, and AMP secretion are all proved to be involved in IBD pathogenesis. As the readout of intestinal barrier function (Fig. 4C), we detected the intestinal permeability evaluated by serum FITC-dextran concentration 4h after oral gavage was significantly enhanced in constipation mice compared to controls (n=4-6, p<0.05). Of note, rhubarb treatment had significantly decreased intestinal permeability (n=4-6, p<0.05). The above results indicated that the intestine barrier was more prone to vulnerability in the constipation group, and more integrity after being administered with rhubarb.

### Results5 Metagenomics analysis

To further reveal the functions and metabolic pathways regulated by constipation and rhubarb treatment, we performed metagenomics analysis. We detected the top 30 bacterial phylum, class, order, family, genes, species in every sample. As the PCA showed (Fig. 5A), the distribution of four groups was significantly different. Figure 5B indicated that constipation induces a significant modification of the diversity and abundance of the gut microbiota composition on the species level, which was perceived as decreased relative abundances of Firmicutes and increased relative abundances of Bacteroidetes in feces. The ratio between these two phyla (the Firmicutes/Bacteroidetes (F/B) ratio) has been associated with maintaining homeostasis, and a decrease in this ratio can lead to bowel inflammation (28). In the constipation group, the percentages of Firmicutes decreased from 16.61% to 26.69% and the percentages of Bacteroidetes increased substantially from 24.4% to 32.76% versus the control group. Surprisingly, the changes partly diminished by treating constipation mice with rhubarb. However, the addition of rhubarb in the normal group imposed little impact on the abundance of Bacteroides and Firmicutes. Similarly, the F/B remarkably decreased after exposure to constipation and increased with the addition of rhubarb. These results may imply that constipation was more prone to inflammation and the tendency was probably reversed by rhubarb treatment.

**Figure 5.**
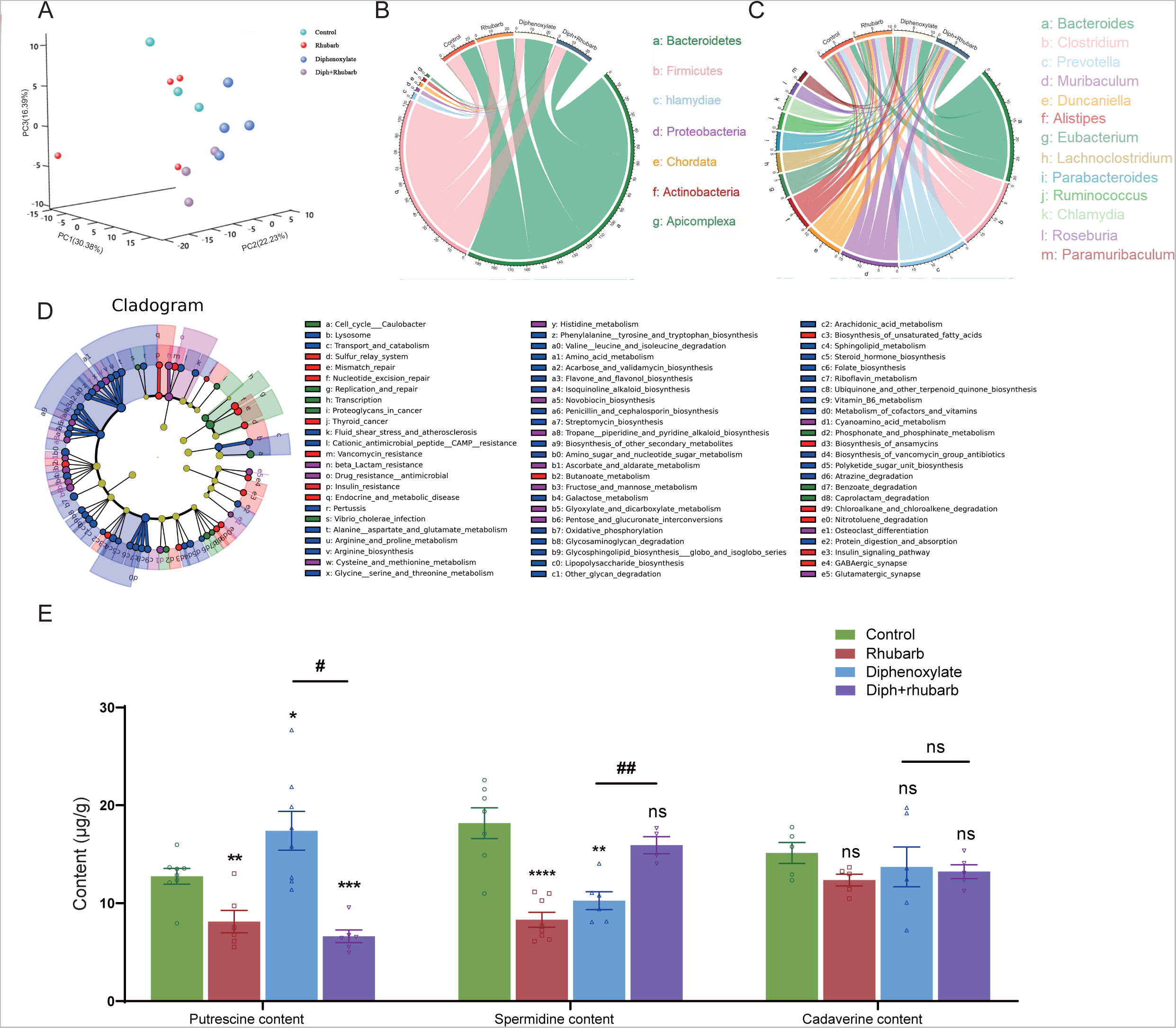
Metagenomics analysis of the feces collected from four groups’ fresh colon and measurements of biogenic amine. (**A**) PCA of 16S metagenomics data of the microbial population in the feces of the four separate groups, each with three to four animals. Comparison of community diversity based on different metric distances of distance metrics useful for microbial communities encoding taxonomic profiles into kernel matrices. (**B**) Circus data reveals the microbiome composition at the phyla level in various groups. (n=3 or 4). (**C**) Circus data shows the composition of microbiota in different groups at the genus level. (n=3 or 4). (**D**) Microbiota functional characterisation based on metabolic pathway abundances. The KEGG BRITE functional hierarchy is represented by a cladogram, with the outermost circles representing individual metabolic modules and the innermost very small circles indicating the KEGG BRITE functional hierarchy. (**E**) The bar graph illustrates that putrescine, spermidine, and cadaverine are measured in the feces (n= 6 or 8). The data is shown as the mean±SEM. ns, P>0.05; *, P<0.05; **, P<0.01; ***, P<0.001. * vs control group; ^#^ vs diphenoxylate group.

Furthermore, we cataloged the genes in the genus level to reve a l differences(Fig. 5C). When rhubarb was administered to normal mice for three days, the levels of Alistipes and Trichinella decreased while Duncaniella, lachnoclostridium, and Parabacteroids increased. While the mice were in constipation, Bacteroides and Muribaculum were enhanced and Clostridium, Roseburia, and Ruminococcus markedly reduced. When the constipation mice were treated with rhubarb, Alistipes, Muribaculum, and Prevotella decreased, and in contrast, Clostridium and Lachnoclostridium increased. This showed that the Diph+rhubarb group was not completely identical to the control group. Alistipes and Ruminococcus were at evidently higher levels than the control group while Bacteroides, Muribaculum, and Parabacteroids were at significantly lower levels.

In addition to relative abundances of microbiota, we detected the abundances of microbial metabolic pathways as profiled from metagenomic shotgun sequencing of a subset of the available body habitats. To identify biological pathways that are regulated by the diversity of the microbiomes, we annotated the genes based on KEGG databases. As for the KEGG databases (Fig. 5D), the gene catalog mainly assigned the top KEGG categories: metabolism, genetic information processing, environmental information processing, cellular process, human diseases, organismal systems and drug development. The results revealed that KOs in the rhubarb group were more abundant which involved in glycolysis/gluconeogenesis (KO00010) and oxidative phosphorylation(K00190) and less abundant in those involved in quorum sensing (KO02024), DNA replication (KO03030) and homologous recombination (KO03440) compare to that in the control group. When constipation, the KOs participate in galactose metabolism (KO00052), oxidative phosphorylation (KO00190) and glycine serine and threonine metabolism (KO00260) were up-regulated, whilst the KOs involved in quorum sensing (KO02024) and mismatch repair (KO03430) were down-regulated. In the Diph+rhubarb group, the KOs representing glycolysis/gluconeogenesis (KO00010), purine metabolism (KO00230), and Aminoacyl-tRNA biosynthesis (KO00970) were less abundant, while the KOs taking part in the quorum sensing (KO02024) and RNA degradation (KO03018) were more abundant compared to the constipation group. However, the up-regulation of functions involved in oxidative phosphorylation, alanine aspartate and glutamate metabolism and biosynthesis of amino acid was remarkable and the down-regulation of functions partaking in DNA replication and mismatch repair was significant. These gene expression changes are statistically significant, with false discovery rates below 0.01.

### Results 6 Treatment of constipation with rhubarb caused changes in biogenic amines

On the basis of the KEGG analysis, amino acid metabolism plays an important role in constipation. Moreover, our present results also indicated that constipation had a significant effect on the gut microbiome and fatty acids (29). It is worth noting that biogenic amine is closely associated with microbiomes. Based on fecal sample analysis, putrescine, spermine, spermidine, and cadaverine are the most common in the human colon (13, 14). Therefore, we investigated putrescine, spermidine, and cadaverine by means of bioinformatics analysis as shown in Figure 5E. It was reported that in vivo and vitro, putrescine impedes intestinal barrier function by disrupting tight junction integrity, aggravates gut leakiness, and subsequently causes disease susceptibility during colonic autoinflammation and infection (30). In line with our expectation, our results indicated that the amount of putrescine increased in the constipation group and dropped to a lesser extent after exposure to rhubarb. Spermidine was reported as being able to take part in maintaining a protective gut microbiota via reducing the expression of genes encoding for a-defensins (DEFAs) by means of transcriptomic and microbiome analyses (31). In contrast to putrescine, spermidine content in the constipation was lower than that in the control group. Likewise, rhubarb contributed to enhancing the abundance of spermidine caused the content to peak in the group treated with rhubarb alone and administration of rhubarb may enable significant reversal of this decreasing effect of constipation on spermidine. Cadaverine, one of a family of small aliphatic nitrogenous bases (polyamines), may be proposed to have the potential to promote bacterial survival under antibiotic exposure and tolerance/resistance formation (32). However, none of these differences were statistically significant among the groups.

### Results7 SCFA and MLCFA

Microbes are metabolically active to survive in the gut environment rather than simply remaining within the gut. Research has pointed out that the intestinal flora has an effect on the progress of composition and numbers of various microbes, the food debris as well as fermentation products such as MLCFAs or SCFAs (33). Gut microbes play an integral role in animal physiology, facilitating metabolism, influencing immunity, and regulating gut function. Numerous studies have confirmed that intestinal flora has an impact on the composition and quantity of various microbes, the food debris as well as fermentation products such as MLCFAs or SCFAs (33). Often, the changes between SCFAs and constipation have been reported (29), but no correlation between MLCFA and SCFA with rhubarb treatment in constipation models. Notably, fatty acid metabolism was involved in the KEGG analysis. To further reveal the association between fatty acids and constipation, we analyzed clustered heatmap drawn based on the Spearman rank correlation matrix, while a hierarchical cluster analysis of all the samples was performed on the correlation coefficients between each pair of fatty acids across all samples (Fig. 6A). The fatty acids, which had the analogous correlations with other fatty acids, were placed close in location. Notably, it can be seen that the four fatty acids (C18.3N3, C18.1N9C, C18.0, C17.0) were clustered into one group, all of which were negatively correlated to another fatty acids group (including C16.1, C18.3N6, C20.1, C20.2, C20.3N3, C20.3N6, C20.4N6, C20.5N3, C21.0, C22.0, C22.1N9, C22.6N3, C23.0, C24.0, C24.1). Next, we performed the correlograms of MLCFA for four groups. There were remarkable differences among the four groups. Constipation caused a significant modification of the interconnections between MLCFA and the median correlation coefficients (0.2097902) (Fig. 6B) was significantly different from the normal group (0.3356643, p < 0.001) (Fig. 6D). In rhubarb group, there were 150 positive correlations decreased, 157 positive correlations increased, 59 negative correlations decreased, 50 negative correlations increased and 89 correlations altered in comparison of the normal group. Compared to the constipation group, there were 92 positive correlations decreased, 163 positive correlations increased, 48 negative correlations decreased, 67 negative correlations increased and 141 correlations altered in constipation. However, there were 83 positive correlations decreased, 165 positive correlations increased, 44 negative correlations decreased, 9 negative correlations increased and 178 correlations altered in constipation. In the constipation group, there were 269 (46.30%) statistically significant correlations (Fig. 6D). In contrast, in normal group, there were 206 (35.46%) such correlations (p< 0.001) (Fig. 6B) and 232 (39.93%) in Diph+rhubarb group (Fig. 6E), which revealed that the MLCFAs in case of constipation were more interactive and recovered when pretreatment rhubarb extract. In addition, no remarkable changes occurred when treated rhubarb extract alone (199, 34.25%). Furthermore, there were strong correlations (|r|>0.75) 103 (17.73%) in normal group, 115 (19.79%) in rhubarb group (ns), 158 (27.19%) in constipation group (p < 0.001) and 184 (31.67%) in Diph+rhubarb group (p < 0.001). It was noted that these results indicated that extremely slight differences were exhibited among the fatty acids that were longer than C20.0 (including C20.1, C20.2, C20.3N3, C20.3N6, C20.4N6, C20.5N3, C21.0, C22.0, C22.1N9, C22.6N3, C23.0, C24.0, C24.1) as shown in Figure. 6B-E. Compared to the normal group (Fig. 6B), the correlations of rhubarb group (Fig. 6C) among the fatty acids that are longer than C8.0 and shorter than C20.1 significantly changed. In line with the changes, the varieties of correlation in the Diph+rhubarb group (Fig. 6E) are also located in the same cites in comparison to those in the constipation group (Fig. 6D), that is to say, the changes of correlation among the C8.0 to C20.0 were remarkable. Strikingly, most of the negative correlations in the constipation group transformed into positive correlations in the Diph+rhubarb group.

**Figure 6.**
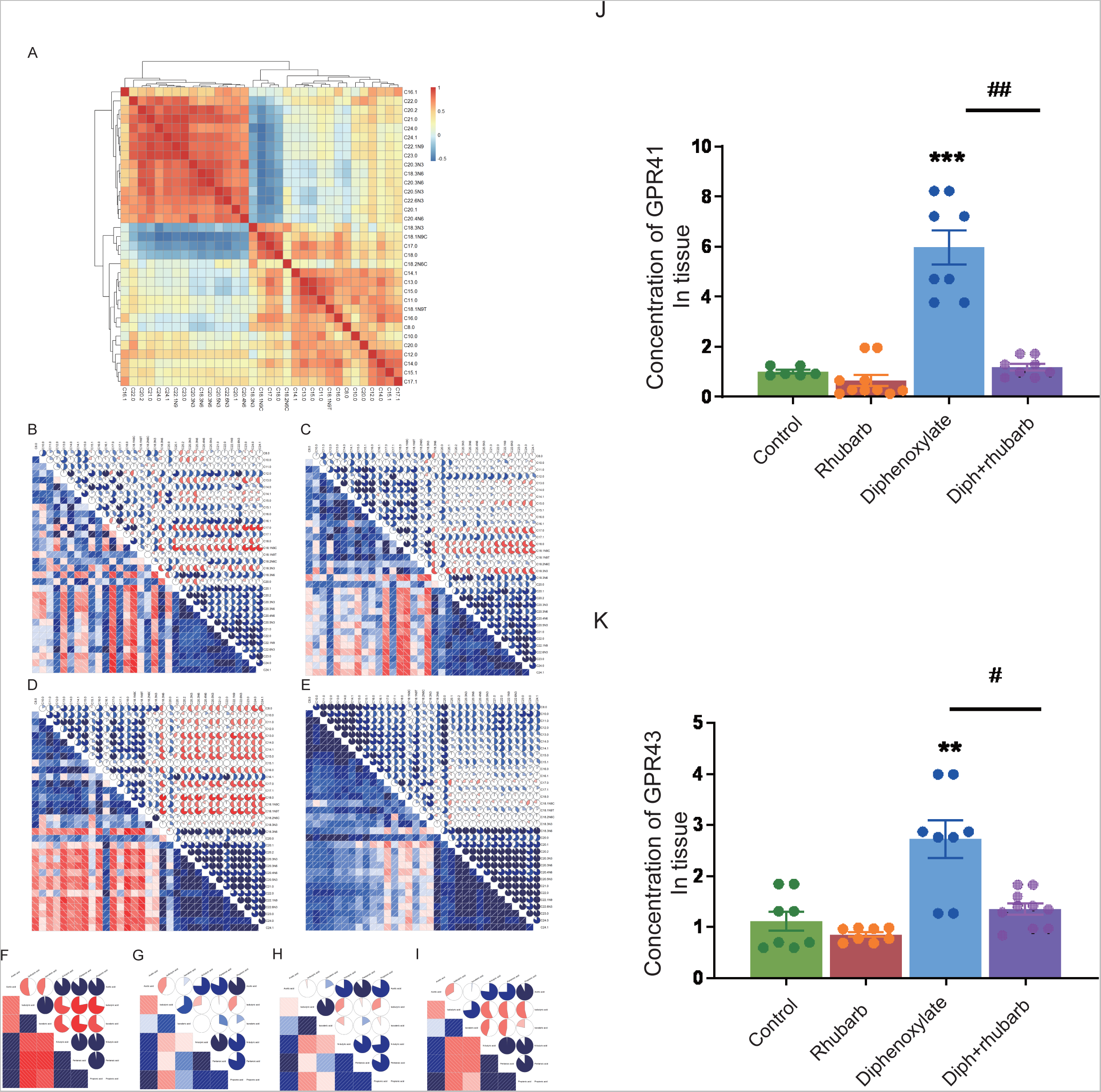
Effect of rhubarb and compound diphenoxylate on the changes of MLCFAs and SCFAs in the feces. (**A**) Spearman rank correlation matrix of the C10-C24 medium-long chain fatty across all samples. The colors are used to denote the correlation coefficients with 1 indicating a perfect positive correlation (red), and -1 indicating a perfect negative correlation (blue). This clustered heatmap was created using the R package “pheatmap.” version 1.0.12. https://CRAN.R-project.org/package=pheatmap. (**B-E**) Correlograms of the C10-C24 medium-long chain fatty in control group (**B**), rhubarb group (**C**), diphenoxylate group (**D**) and Diph+rhubarb group(**E**). (**F-I**) Correlograms of matrix of the short chain fatty in normal(**F**), rhubarb(**G**), constipation(**H**) and Diph+rhubarb group (**I**). (**J-K**) The bar graph illustrates the concentrations of SCFA receptors, GPR41/ GPR43 from the four groups. The data is shown as the mean±SEM. ns, P>0.05; *, P<0.05; **, P<0.01; ***, P<0.001. * vs control group; # vs diphenoxylate group.

Finally, we disclosed the correlograms of SCFA in four groups. As Figure. 6F-I showed, 19 (90.48%) statistically significant correlations among SCFAs existed in normal groups and 13 (61.90%) such correlations in the other three groups. As indicated in Fig. 6G, for example, the correlations of isovaleric acid in the rhubarb group had an obvious difference compared to those in the normal group (Fig. 6F). The same conclusion was reached when the Diph+rhubarb group shown in Fig. 8I was compared to the constipation group (Fig. 6H). The above results demonstrated the fatty acids were affected by constipation and rhubarb.

### Results8 Changes in the expression of GPR41 and GPR43 in different groups

SCFAs may signal through cell surfaces, like GPR41, GPR43, and GPR109A, to activate signaling cascades and play a pivotal role in perpetuating intestinal inflammation. So we evaluated the expression of GPR41 and GPR43 in the colon tissue (Fig. 6J and K). As the figure shows, their expressions both increased significantly in the constipation group, while no differences were shown in the Rhubarb group or in the Diph+rhubarb group compared to the control group. It’s noteworthy to state that their high expressions in the constipation group were rectified after being treated with rhubarb, which elucidated that rhubarb may have the suppressive inflammatory effect.

## 3. Discussion

Rhubarb is an effective Chinese herb used to relieve constipation that has aroused much attention due to its great amount of usage. To unveil the mechanism of relieving constipation by rhubarb, we generated the constipation model. In this study, our data suggested that the muscle layer and the new collagen in constipation were significantly increased, which displayed that constipation may induce fibrosis. Of note, rhubarb has the ability to reverse the stiffness to recover muscle strength with the symbol of a thinner muscle layer and reduced collagen. Moreover, our previous studies have revealed that promoting colonic mucus synthesis and secretion. Colonic mucus secreted from goblet cells is attached to the epithelium and isolates the epithelium from external environment (34). The results above verify that rhubarb may relieve constipation by strengthening muscles and boosting mucin secretion to promote defecation.

Our data revealed that the constipation mice with the decreased mucus layer were prone to have the impaired barrier and increased permeability with the manifestation of a high concentration of FITC-dextran. The condition also created an opportunity for bacteria to invade and induce inflammation, which accounted for the increased pro-inflammatory cytokines, such as IL-17A and IL-23. It is worth noting that the tendency of IL-22 was diverted as it peaked in the rhubarb group and fell in the constipation group. As the previous experiments demonstrated that IL-22 could work as a contributor to maintaining the mucus barrier function (27), the current result was in accordance with our speculation that rhubarb contributed to maintaining the tightness and integrity of the intestinal barrier.

MyD88, a fundamental role in the innate immune system, is the primary adaptor protein not only of IL-1 and IL-18 receptors but also of almost all the TLRs and thus considered as a central hub of the inflammatory signaling cascades as well as is found to be required in LPS signalling. An interesting report concluded that IL22 induced significant upregulation of transcripts involved in microbial sensing (Tlr4, Myd88, Tnfaip3) (35). NF-κB, activated by TLR stimulation, is a key regulator of inflammation, innate immunity, and tissue integrity (36, 37). The tendency of LPS, NF-κB, TLR4 and MyD88 were coincident in the four groups. The high expressions after being treated with rhubarb were uncommon due to their pro-inflammatory role, however, there are several reasons that may make sense. Firstly, rhubarb treatment arouses the activation of the mast cells, which plays an important role in immunity, as our previous report examined. On the other hand, the plasma cells were accumulated and activated, which is a critical step for the innate immunity. Moreover, TLR/NOD ligands have been shown to modulate mucin gene expression and promote mucin secretion from goblet cells (38), which may be one of the mechanisms that rhubarb promote mucin secretion.

What role does intestinal flora play in this process? Intestinal flora contains about 1000 different bacteria (39) and has a sophisticated effect on immunity and metabolism particularly (40). It was emphasized that lots of diseases have demonstrated altered bacterial diversity comes along with reductions in the abundance of beneficial microorganisms due to the fragility of the gut microbiota (41). Imbalances in the gut microbiota result in a basal inflammatory state and enhanced susceptibility to viral and bacterial infections (42). According to the PCA result, the number of gut microbial species, bacterial abundance, and flora diversity were remarkably different in the different mice models. Specifically, the exposure to constipation enabled the potential of decreasing the diversity of the microbiome and characterized the high concentration of Bacteroidetes and markedly decreased the F/B ratio. A stream of a pilot study reported the predisposition to inflammation sensitivity caused by the decreased F/B ratio. Microbes are metabolically active to survive in that gut environment rather than simply remaining within the gut. Hence, the intestinal flora would play an important role in many areas, for example, the progress of composition and numbers of various microbes, food debris, and fermentation products. To gain functional insights into colon metabolism, we assessed the genes by KEGG analysis. According to the variance analysis, it captured the preference for the amino acid metabolism. In addition, glycometabolism was also involved in the process. Therefore, we applied the MS analysis to determine the alteration of biogenic amine and fatty acid. Polyamines have attracted much interest, in part, because of their essential roles in multiple cellular functions, like cell growth, mitochondrial metabolism and histone regulation (17, 43–48). Gut microbiota can produce the bacterial biogenic amines, including putrescine, cadaverine, tyramine, and 5-aminovalerate from amino acid degradation (arginine, lysine, tyrosine, and proline, respectively). It is beyond all doubt that our data elucidate that putrescine and spermidine significantly changed. It was later found that MLCFA and SCFA were altered. SCFA has an impact on maintaining homeostasis in the colon and supplies 60%–70% of energy that colonic epithelia need (33). SCFAs can be produced by bacteria. Notably, the number of bacteria, the pH and the substrate can notably influence the process (49). Previous studies have shown that different substrates produce different amounts and proportions of SCFAs, which participate in many critical physiological metabolic processes in vivo such as induction of cell differentiation, regulation of the growth and proliferation of normal colonic mucosa and reduction of the growth rate of colorectal cancer cells. As our investigation shows, the composition and diversity of the intestinal flora have varied . It can be seen that SCFAs, such as butyrate including N-butyrate and isobutyrate, pentanoic acid, isovaleric acid, increased significantly after rhubarb administration, but decreased significantly in the constipation group in our research. N-butyrate and pentanoic acid have a more significant decrease further in the treatment group, while isovaleric acid increased significantly. On the contrary, much of the research on this topic demonstrated that the content of isobutyrate in samples from subjects with constipation is significantly higher than in those from healthy people (50). The diet might make sense of the phenomena, given that SCFAs originate from the degradation of polysaccharides. Emerging evidence ha s come to suggest that SCFAs and MCFAs were mainly esterified by long-chain fatty acid groups, and SCFA and MCFA concentrations in full-term milk were significantly higher than those in premature milk (51). The correlation between SCFAs and MLCFAs in feces, especially in the alteration of intestinal flora, needs further study. Emerging studies highlight the importance of SCFAs in activating GPR41 and GPR43 on intestinal epithelial cells, resulting in mitogen-activated protein kinase signaling and rapid production of chemokines and cytokines (13). These pathways regulate protective immunity and tissue inflammation in mice. High concentrations of GPR41 and GPR43 in constipation mice strongly show perfect concordance with the results directing inflammation in constipation

## 4. Conclusion

Collectively, the most obvious finding to emerge from this study is that constipation was linked to inflammatory response and gut microbiota as well as metabolic disorders. Notably, rhubarb treatment may play the regulatory and reversing role in these biological process es to relieve constipation through a multitude approach. Undeniably, the major limitation of this study is that Rhubarb was used in this experiment rather than its active ingredient, which resulted in a complicated effect. Notwithstanding these limitations, the study suggests that this new work should therefore assist in our understanding the role of rhubarb as a new multi-target drug for clinical application.

## 5. Materials and methods

### Animals

C57B/6N male mice weighing 21±1g from Laboratory Animal Services Center of Capital Medical University are raised under standard environment (22.0-25.0 ℃, at a relative humidity of 50–70% under 12-/12-hour light/ dark cycle) and all procedures were carried out according to National Institutes of Health Guide for the Care and Use of Laboratory Animals (AEEI-2016-079).

### Regents and Dosage Information

Compound diphenoxylate containing 25 mg of diphenoxylate and 2.5mg of atropine sulfate monohydrate per tablet was purchased from Hefeng Medicine Industry (Guangxi, China, Lot: 210704). The compound diphenoxylate was dissolved in normal saline to achieve an adequate concentration of 10 mg/ml. Administration of compound diphenoxylate to mice at the dose of 20 mg/kg via gavage lasting five days was prepared as a verifiable and repeatable constipation model group (52, 53) as the constipation model group. We purchased rhubarb from Tongrentang Pharmacy (Beijing, China) and identified it with the assistance of Prof. W. Wang from Xuanwu hospital, Capital Medical University. As described previously (54), the roots of therhubarb were crushed and soaked in the annealing for 2h and stored at 4°C until use. Previous experiments studying strongly driven systems have reported remarkable effects at doses of 600mg/25g. This is important because the optimal dose elicits alleviation without any other side reaction. Additionally, in an analysis of a large randomized clinical trial of constipation, the dose application was judged to be about 9 fold to those administered in human clinical trials adult dosage (55).

### Experimental design

The mice were randomly divided into four groups with the same number of animals in every group: the control group received normal saline alone on the same day as the other groups. Another group of mice was treated with normal saline vehicles once daily for five days, then with three-day rhubarb extract followed at the dose of 600mg/25g. To induce constipation, mice undergoing administration diphenoxylate for five consecutive days were separated into two groups, one with three-day normal saline treatment, one co-administration rhubarb extract for three days. All mice were raised in the metabolic cage with free access to food and water to collect 24-h feces and supervised the consumption of food and water so as to accurately judge the success of model mice (56–58). The detailed information on experimental design is available in exhibited Figure 1A.

At the end of the experiment, feces from the colon and ileocecus, urine from bladder and blood were collected before euthanasia. At the end of the sacrifice, the colon was collected and dissected out to cut open for measuring colon length and other downstream analysis. The samples of blood were collected then centrifuged at 12000 r/min for 30 min at 4℃ in order to obtain the serum and stored at -80℃. The feces were stored in the sterile centrifuge tubes at -80℃ until being performed 16S rDNA and further metagenomic analysis. The colon tissue was removed from the point of 0.5cm above the anus to top of ileoceca and soaked in different fixative solutions. The tissue pieces fixed in 10% formalin overnight at room temperature were stained with Masson trichrome, Sirius Red or hematoxylin and eosin for interstitial image datasets with light microscopy. The samples were paraffin-embedded, and 5-µm-thick serially sections were mounted on glass slides and stored at -20℃. Paraffin sections were first dewaxed and rehydrated through a graded alcohol series and transparently with xylene before antigen retrieval.

### Histological studies

#### HE staining

The sections were stained in parallel employing by modified Lillie-Mayer’s hematoxylin for 1 min, differentiated with 1% hydrochloric acid alcohol for 2-5 seconds, and soaked in tap water for 10 min to make it blue followed by dyed with water soluble red dye for 1 min where indicated.

#### Masson trichrome staining

To evaluate whether collagen fibers are changed during this process, Masson’s trichrome staining was performed with a commercial kit (Beijing Solarbio Science & Technology Co.,Ltd). The sections were stained using Wiegert’s iron hematoxylin solution (Wiegert solution A: Wiegert solution B =1;1) for 10 min and then stained with azaleine for 10 min at room temperature, respectively followed by weak acid solution (deionized water: weak acid =2:1) washed along with immersing phosphomolybdic acid for 2 min and stained in diluted toluidine blue for 1 min. All slices were washed five times with weak acid solution until the collagen fibers to the total area appeared as blue. Similarly, the collagen fiber densities and distribution were quantified with Image-pro plus.

### Sirius Red

Sirius red staining, as well as Masson trichrome stain (MTS), was detected for histologically assessing collagen content followed by a combination of microscopic including polarized light and optical microscopy. The sections were rehydrated and stained for 1 h with a Sirius red stain kit (0.1% Sirius red in a saturated aqueous solution of picric acid) (Beijing Leagene Biotechnology Co.,Ltd.)

### Atomic force microscopy(AFM)

The muscle layer variation was observed in the response of different treatments and we wonder whether the muscle change along with the strength change has an important role to play. Therefore, AFM was implemented using a Multimode/Nanoscope IIIa AFM (Digital Instruments/Veeco, Santa Barbara, CA). MLCY-BIO (BRUKER, USA) with a nominal spring constant of 0.14 N/m, which is capable of detecting samples with regard to its stiffness, adhesion, and modulus. The colon specimen was removed immediately, embedded in OCT, and snapped frozen in liquid nitrogen and substantially restored at -80 ℃ . The colon tissue slices were sliced into 5μm sections. The pieces were fixed with 4% PFA for 30 min and washed with PBS mixed with 1% cocktail (27423400, Switzerland) for 5 min for a total of three times. These tissues went through imaging under the AFM imaging system. All images were detected in the intermittent contact mode in regime liquid at room temperature.

### Enzyme-linked immunosorbent assay

Levels of cytokines in the serum of all the mice (TNF-a, IFN-γ, IL-1β, IL-6, IL-12, IL-17, IL-4 and IL-10) were performed using commercially available mouse ELISA kits according to the protocols supplied by the manufacturer and detected by a multimode microplate reader (Beckmancoulter UniCel DxC 600 Synchron, U.S.). The total protein in each sample was measured by TP Kit RGB& CHN(Lot:20221218.30002).

### In vivo Paracellular Permeability Assay

In order to assess colonic paracellular permeability in vivo, mice were deprived of food for 18 hours, then orally gavaged with 440 mg/kg body weight of FITC-labeled dextran (FD4) (Sigma, St. Louis, MO, USA). The mice were sacrificed 4 hours later, and plasma was collected and its fluorescence intensity in serum was detected by a fluorescent microplate reader (excitation at 480 nm and emission at 520 nm; HTX Multi-Mode reader, SYNERG).

### Real-Time PCR Analysis

Total RNA was extracted from prepared tissue using FastPure Cell/Tissue Total RNA Isolation Kit V2 (RC112, Vazyme, Nanjing, China) according to the product manual. The concentrations of isolated RNA were quantified by NanoDrop 2000 spectrophotometer (Thermo Fisher Scientific, Waltham, MA), and then reverse transcription was performed by HiScript III RT SuperMix for qPCR kit (R323, Vazyme, Nanjing, China) by BIO-RAD iCycler(BIO-RAD, USA). Finally, the cDNA was with Taq Pro Universal SYBR qPCR Master Mix kit (Q712, Vazyme, Nanjing, China) by the CFX96TM Real-Time System (BIO-RAD, USA). The thermal cycles were 95 ℃ for 5 min, 56℃ for 15 min, 72℃ for 10 minutes, for 45 cycles, and 60℃ for 1 min. The relative amount of the target mRNA was normalized to the GAPDH level, and data were calculated by the 2^−ΔΔCt^ method. The primer sequences were listed as follows.

GPR41: forward CTTCTTTCTTGGCAATTACTGGC;

reverse CCGAAATGGTCAGGTTTAGCAA.

GPR43: forward CTTGATCCTCACGGCCTACAT;

reverse CCAGGGTCAGATTAAGCAGGAG.

GAPDH: forward AGTGTTTCCTCGTCCCGTA;

reverse CGTGAGTGGAGTCATACTGG.

### LC-MS / MS Metabolite analysis

Targeted feces metabolomics quantifying fatty acids were performed by LC-MS/MS processes previously reported by Gao *et al*. (29). Fecal samples were briefly homogenized in a Bullet Blender into suspension, then hydrochloric acid (30mM) was added, isotopically-labeled acetate (0.125 mM), butyrate hexanoate (0.125 mM) and 250 mL of Methyl tert-butyl ether (MTBE). Finally, each sample is a mixture of 400 ml in volume. Subsequently, the mixture was briefly mixed by vortexing for 10s at 4°C twice and the solvent layers were separated by centrifugation for 1 min. 10 mL of MTBE epurated from the samples was laterally transferred to an autosampler vial for GC-MS analysis by placing it in a separate auto-sampler vial to get a series of calibration standards for normal quality purposes. GC-MS analysis of samples was implemented with Agilent 69890N GC-5973 MS detector with the parameters given in extended methods. A 1μL sample was injected with a 1:10 split ratio on a ZB-WAXplus, 30m 0.25 mm x 0.25μm (Phenomenex Cat# 7HG-G013-11) GC column. Helium was used as the carrier gas at a flow rate of 1.1 mL/min with 240°C as the injector temperature, and the column temperature was kept at 310°C under isocratic condition. Quantification data were extracted and analyzed by MassHunter Quantitative analysis version B.07.00. SCFAs were normalized to the nearest isotope labeled internal standard and quantitated using 2 replicated injections of 5 standards to create a linear calibration curve with accuracy greater than 80% for each standard (59).

### DNA sequencing and metagenomic sequencing

Collected stool samples were collected in sterile tubes and immediately frozen as well as kept at -80°C until performing further analysis. DNA was extracted with HiPure Stool DNA Kit (Shanghai Ponsure Biotech, China) and measured concentration and quality. Quantitative real-time PCR was performed using bacterial primers which were targeted amplification of the combined V3 and V4 regions of the 16S rDNA gene. Amplification was performed with fusion primers containing the 16S-only V3-V4 sequences fused to Illumina adapters overhung nucleotide sequences (60) and finally pooled and sequenced on Illumina’s MiSeq/NovaSeq platform at the Genomic and Proteomic Core Laboratory in Genewiz, LTD, Suzhou, China. The generated NGS data was filtered and clustered into operational taxonomic units (OTUs), which carry species distribution information. Based on the above data, a series of analysis methods was employed to unveil the difference among multiple groups.

The metagenomic DNA in the colon content of mice in each group was executed using the Stool Genomic DNA Kit (CoWin Biosciences, China) and used for quantitative analysis of gut microflora. Then, the purified DNA was end-repaired using the End-it End-repair kit and then added an “A” base to the 3’end of DNA fragments. Additionally, for adaptor ligation, paired-indexed Illumina dual end adapters were replaced with palindromic forked adapters with unique 8-base index sequences embedded within the adapter and added to each end. Target DNA fragments within a certain range of length were screened by the magnetic beads, and amplified with PCR with index at the end of the target fragment to complete the construction and detection of the sequencing library. We prepared sequencing libraries using Illumina’s TruSeq ChIP Library Preparation kit, and barcoded libraries on an Illumina HiSeq2000 instrument according to the fragment size. Lastly, we generated gene profiles using gene catalogue and estimated these data by the data library KEGG (Kyoto Encyclopedia of Genes and Genomes) ortholog (www.genome.jp/kegg/).

## 6. Statistical analysis

All data other than the sequencing data were plotted and analyzed with Prism Software 8.0 (GraphPad Software, San Diego) and presented as mean ± standard error of mean (SEM). Comparisons between groups were performed using ANOVA with post-hoc tests or Student’s t-tests. A p-value less than 0.05 was considered statistically significant, and one between 0.05-0.10 as showed a trend toward statistical significance. Principal component analysis (PCA) based on the unweighted UniFrac distance metrics was used to assess Beta diversity. Pearson r coefficients were applied to calculated bivariate correlations and paired Mann-Whitney’ test was used to compare p-values between groups. Correlation matrices also were displayed as schematic correlograms (61). All statistical analyses were performed in Stata/SE 12 and open source procedure R 4.1.1 (https://www.r-project.org/).

## 7. Study approval

The study was approved by Animal Care and Use Committee of Capital Medical University (AEEI-2016-079).

## 8. Author contributions

Han Gao generated the mice model. Han Gao, Chengwei He, Rongxuan Hua, Chen Liang and Yixuan Du performed experments. Lei Gao conducted the bioinformatics analysis. Hongwei Shang performed the experiments and supplied the experimental instructions. Rongxuan Hua, Yuexin Guo and Lucia Zhang analyzed the data from the experiments. Boya Wang and Lucia Zhang drew the graph and expanded the literature. Jingdong Xu cooperated, analyzed all the data and revised the manuscript.

## 9. Dataset availability

The data in this study generated during and/or analyzed during the current study and supplementary information are obtained from the corresponding author on reasonable request.

## Supporting information

graphic abstract

## Acknowledgements

This work was supported by the National Natural Science Foundation of China Grant (No.82174056, 81673671 JD Xu).

## Notes

Conflict-of-interest statement: The authors have declared that no conflict of interest exists.

### Competing Interest Statement

The authors have declared no competing interest.

