## Supplementary figures and images for "Underlying beneficial effects of Rhubarb on constipation-induced inflammation, disorder of gut microbiome and metabolism"

### graphic abstract

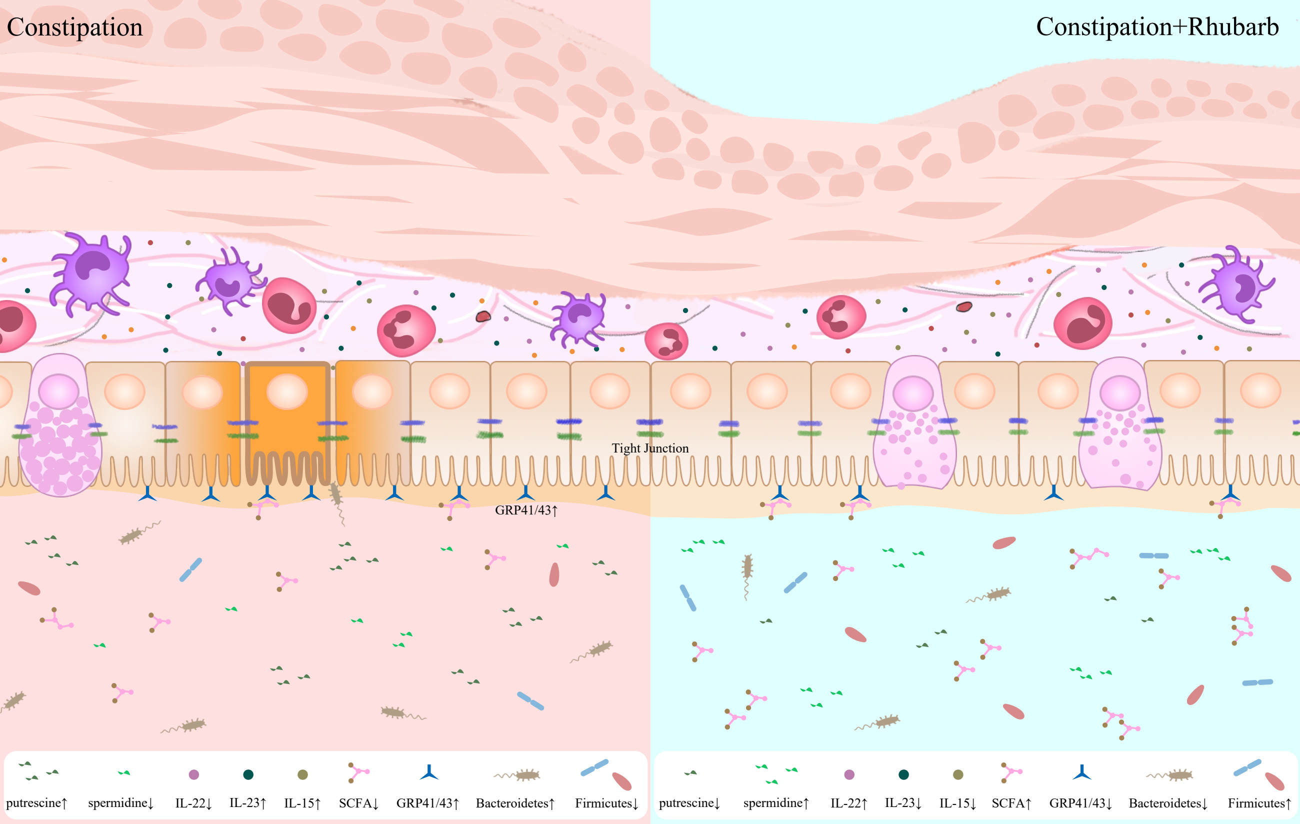
